# Combinatorial approach of binary colloidal crystals (BCCs) and CRISPR activation to improve induced pluripotent stem cell differentiation into neurons

**DOI:** 10.1101/2020.12.21.423852

**Authors:** Daniel Urrutia-Cabrera, Roxanne Hsiang-Chi Liou, Jiao Lin, Kun Liu, Sandy S.C. Hung, Alex W. Hewitt, Peng-Yuan Wang, Raymond Ching-Bong Wong

## Abstract

Conventional methods of neuronal differentiation for human induced pluripotent stem cells (iPSCs) are tedious and complicated, involving multi-stage protocols with complex cocktails of growth factors and small molecules. Artificial extracellular matrix with defined surface topography and chemistry represents a promising venue to improve the neuronal differentiation *in vitro*. In the present study, we test the impact of a type of colloidal self-assembled patterns called binary colloidal crystals (BCCs) in neuronal differentiation. We developed a CRISPR activation (CRISPRa) iPSC platform that constitutively expresses the dCas9-VPR system, which allows robust activation of the proneural transcription factor *NEUROD1* to rapidly induce neuronal differentiation within seven days. We showed that the combinatorial use of BCCs can further improve this neuronal differentiation system. In particular, our results indicate that fine tuning of silica and polystyrene size is critical to generate specific topographies to improve neuronal differentiation and branching. BCCs with 5 μm silica and 100 nm carboxylated polystyrene has the most prominent effect on increasing neurite outgrowth and more complex ramification, while BCCs with 2μm silica and 65nm carboxylated polystyrene is better in promoting neuronal enrichment. These results indicate that biophysical cues can support rapid differentiation and improve neuronal maturation. In summary, our combinatorial approach of CRISPRa and BCCs provides a robust and rapid pipeline for *in vitro* production of human neurons. Specific BCCs can be adapted to late stages of neuronal differentiation protocols to improve neuronal maturation, which have important implications in tissue engineering, *in vitro* biological studies and disease modeling.

## Introduction

Neurons in the central nervous system form a complex cellular network that is critical for the transmission and processing of signals from the body and its surroundings. However, neurons have limited regenerative capacity, as such neurodegeneration caused by trauma or disease often results in permanent damages to the nervous system ^1^. Furthermore, the limited availability of *bona fide* human neurons to use for *in vitro* studies, represents a major challenge hindering the study of neurodegenerative diseases and development of cell replacement therapies.

Advances in induced pluripotent stem cell (iPSC) technology provided a cellular source that gives us unprecedented access to nearly all neuronal subtypes. The derived neurons can be employed to develop *in vitro* disease models for drug discovery, or development of cell therapy to treat neurodegenerative disorders ^2–4^. Conventional neuronal differentiation protocols often recapitulate the signalings during neural development using various cocktails of growth factors and small molecules. However, many of these neuronal differentiation protocols yield variable results and are difficult to upscale, as they require long-term culture using media with complex compositions ^2,5–7^. In addition, neuronal maturation is difficult to achieve as many differentiation protocols yield heterogeneous populations with varying maturation stages ^6,8,9^. Therefore, there is a need to improve differentiation protocols to generate enriched neuronal populations with mature characteristics, while simplifying the cumbersome culture conditions.

Transcription factor-mediated differentiation is a promising approach to promote iPSC differentiation into neurons. Notably, overexpression of key proneural transcription factors can drive neuronal differentiation with considerably shorter time compared to growth factors ^7,10,11^. However, further optimisation is needed for *in vitro* maturation of neurons, such as long neurites and complex branching which facilitates formation of neuronal networks ^2,12^. A common strategy to promote neuronal maturation is to prolong the *in vitro* culture, which typically adds many weeks to the protocol and significantly increases the cost of *in vitro* production of neurons.

It is now clear that cell-matrix interactions play an important role in modulating cellular behaviour, which could be harnessed to improve iPSC differentiation into neurons. Indeed, many groups have shown that the substrate topography can influence proliferation, morphology, and gene expression ^13–16^. In particular, topographical patterns like gratings and pillars have been employed to support the generation of neurons derived from stem cells, ^13,17–19^ or neurons derived from fibroblasts using direct reprogramming ^20,21^. Our group has previously developed topographically and chemically defined surfaces that are suitable for stem cell culture ^15,22^, which are based on binary colloidal crystals (BCCs) ^23^. We also showed that BCCs can assist in iPSC reprogramming protocols by increasing the proportion of fully reprogrammed human iPSC colonies ^22^. Using a small molecule approach, BCCs improve the cardiac differentiation of human iPSCs via morphological and biological manipulation ^24^.

In this study, we explored the effect of BCCs as an artificial extracellular matrix for neuronal differentiation. We developed an iPSC platform for CRISPR activation (CRISPRa), which allows efficient activation of endogenous genes ^25^. Using this system, we showed that the sole activation of *NEUROD1* allows rapid generation of neurons within 7 days. Interestingly, the combinatorial use of BCC further promotes neuronal enrichment and maturation with increased neurite outgrowth and complexity. Our study demonstrates a novel approach using topographical cues such as BCC together with CRISPR technology to improve the differentiation and maturation processes of stem cell-derived neurons.

## Results

### Characterisation of the binary colloidal crystals (BCCs)

From a library of BCC that we generated ^32^, we selected three uncharacterised BCC monolayer surfaces with different properties of roughness and wettability. The three BCC surfaces have different topographies consisting of varying sizes of Si particles and polystyrene with or without carboxylation (Figure 1A): BCC9 (5 μm Si particles and 100 nm carboxylated polystyrene), BCC13 (2 μm Si particles and 65 nm polystyrene) and BCC16 (2μm Si particles and 65 nm carboxylated polystyrene).

**Figure 1:**
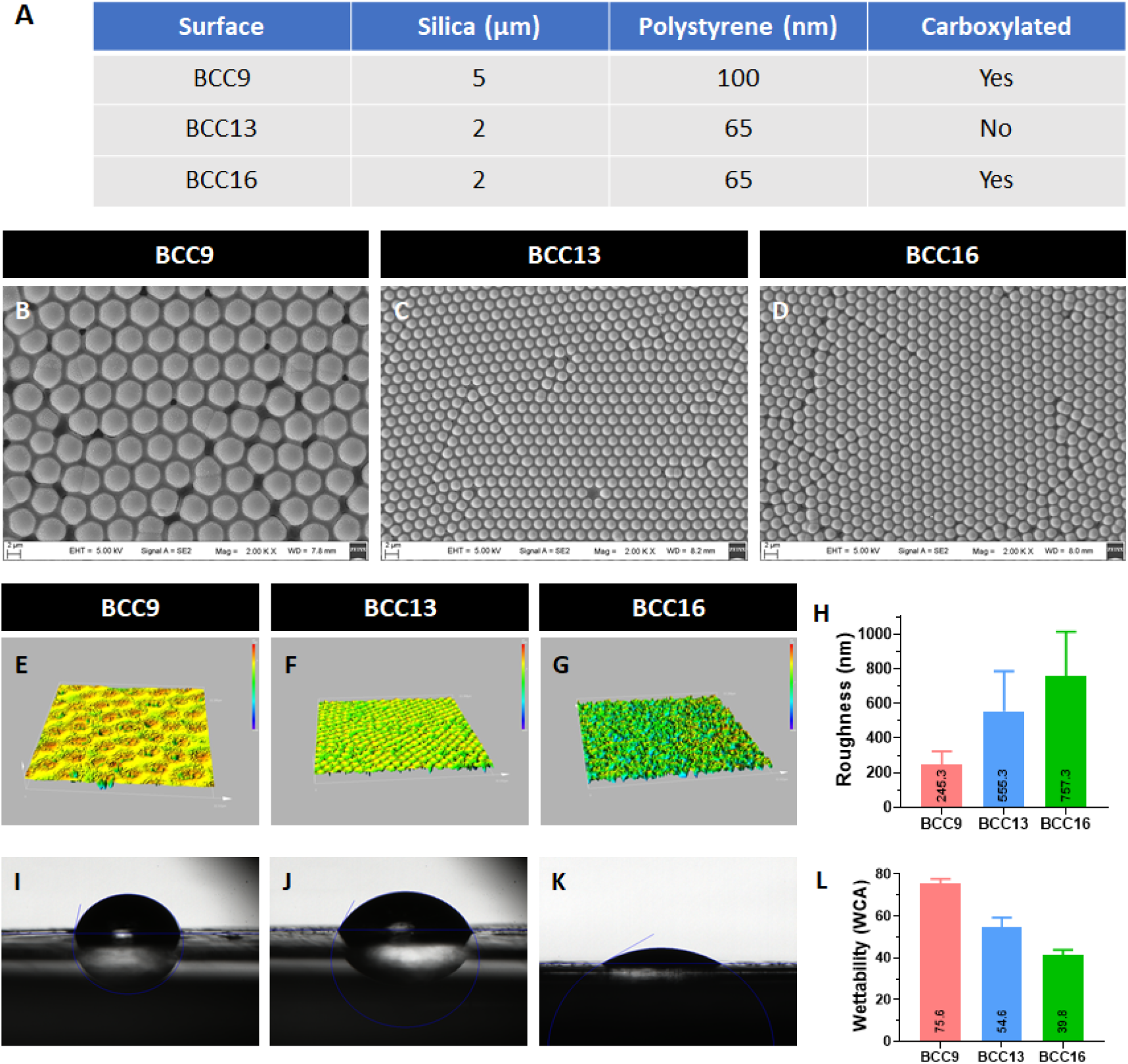
Characterisation of binary colloidal crystal (BCC) surfaces. A) Composition and properties of the three BCC surfaces, BCC9, BCC13 and BCC16. Analysis of the pattern and structure of BCCs using field emission scanning electron microscopy for B) BCC9, C) BCC13 and D) BCC16. E-G) Surface roughness analysis for the three surfaces and H) the pooled quantification (n=3, error bars represent SEM). I-K) Wettability analysis showing the water contact angle (WCA) for the three surfaces and L) the pooled quantification (n=3, error bars represent SEM).

Detailed characterization of the three BCC surfaces was performed. Scanning electron microscopy showed that the three BCCs have high quality ordered surface symmetry and defined topography and are close packed as hexagonal structures (Figure 1B-D). BCC9 is the smoothest surface among the three (roughness = 245.3nm), followed by BCC13 (555.3nm) and BCC16 (757.3nm, Figure 1E-H). Notably, BCC13 and BCC16 have the same Si particle size but different surface roughness, indicating that carboxylation of polystyrene increased surface roughness via self-assembly (Figure 1H). Also, our analysis for surface wettability showed that BCC9 was the most hydrophobic (mean WCA =75.6 degrees), followed by BCC13 (mean WCA = 54.6 degrees) and BCC16 was the most hydrophilic surface (mean WCA = 39.8 degrees, Figure 1I-L). Overall, the three BCCs have varying degrees of surface roughness and wettability similar to those we used for iPSC culture ^22^, with BCC9 being the smoothest and most hydrophobic and BCC16 being the roughness and most hydrophilic. This allows us to study the effect of different topographies in manipulating neuronal differentiation.

### Generation of human iPSC from PBMC

To study the effect of BCCs in neuronal differentiation, we first generated an iPSC line using peripheral blood mononuclear cells (PBMC) obtained from a healthy 65 year old male (Figure 2A). The derived iPSC line, named PBMC-iPSC, was carefully characterised to confirm pluripotency and its quality. Immunocytochemical analysis showed that the iPSC expressed the pluripotency markers OCT3/4 and TRA-1-60 (Figure 2B-C). Using embryoid bodies assay (Figure 2D, we showed that PBMC-iPSC retains the potential to differentiate into the three germ layers *in vitro*, as demonstrated by expression of βIII-Tubulin (ectoderm), smooth muscle actin (SMA; mesoderm) and alpha-fetoprotein (AFP; endoderm, Figure 2E-G). Furthermore, we confirmed that PBMC-iPSC maintains a normal karyotype using a copy number variation assay (Figure 2H). Together, these characterisation results confirm the quality of the derived PBMC-iPSC line.

**Figure 2.**
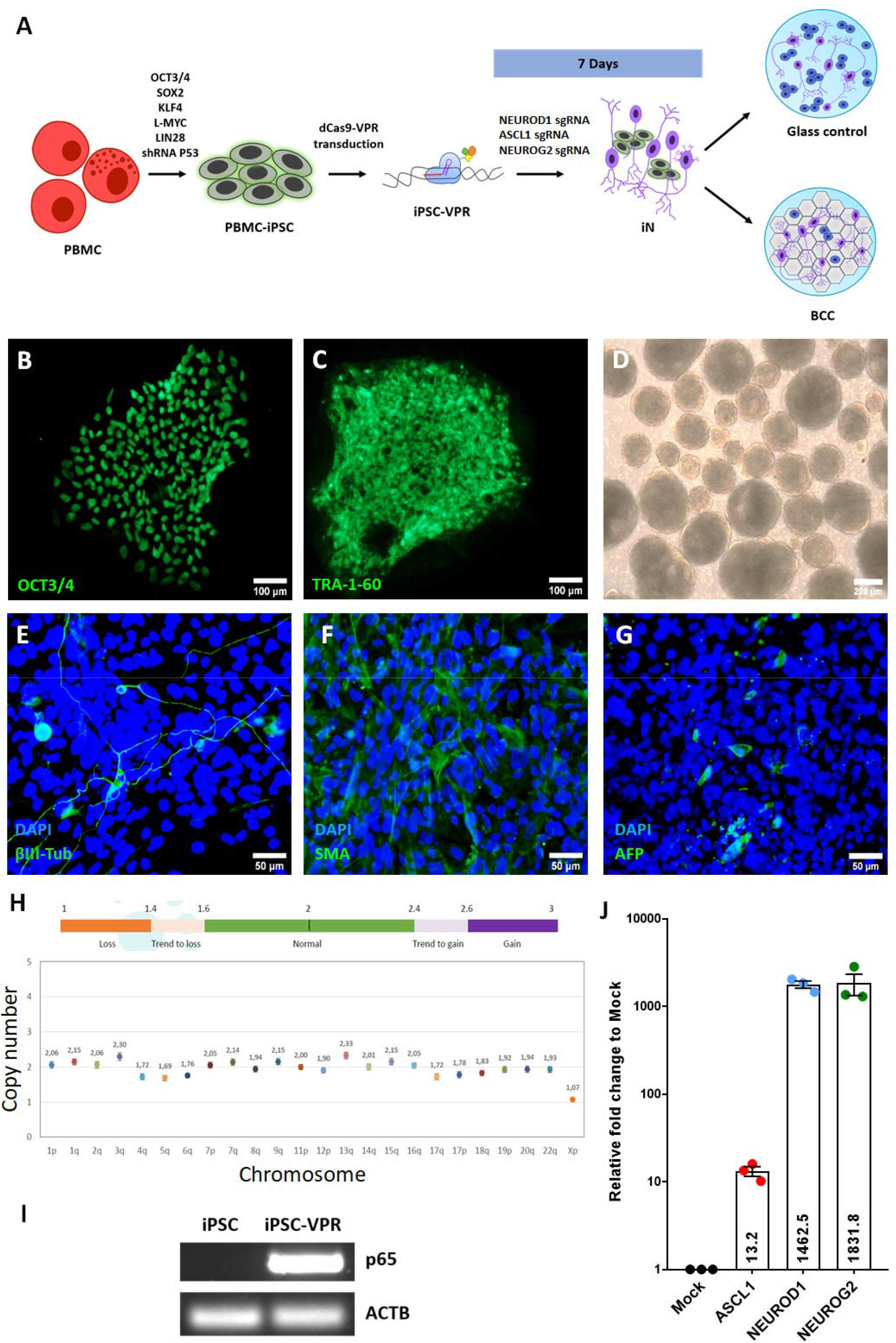
Generation of a CRISPRa iPSC system: A) Schematic representation of our study. Immunohistochemistry analysis of the pluripotency markers B) OCT3/4 and C) TRA-1-60 in PBMC-iPSC. Scale bar = 100 μm. D) Embryoid bodies formation yield differentiated cells representative of the three germ layers, as indicated by marker expression of E) βIII-tubulin (ectoderm), F) SMA (mesoderm) and G) AFP (endoderm). Scale bar = 50 μm. H) Copy number variation analysis of the karyotype of PBMC-iPSC. I) RT-PCR analysis of the p65 component of VPR activator in parental PBMC-iPSC (iPSC) and the stable iPSC line expressing dCas9-VPR (iPSC-VPR). J) qPCR analysis of CRISPR activation for the proneural factors *ASCL1, NEUROD1* and *NEUROG2* in iPSC-VPR. n= 3 biological repeats, error bars represent SEM.

### Transcription factor-mediated neuronal differentiation using CRISPRa

Transcription factor-mediated differentiation has been demonstrated to be a rapid approach to induce differentiation of pluripotent stem cells into neurons ^7^. Previous reports have shown that transgene overexpression of three master regulators for neuronal specification, *ASCL1, NEUROD1* or *NEUROG2*, are able to direct iPSC to differentiate into neurons ^10,25,33^. To establish a robust platform for neuronal differentiation, we utilised the CRISPRa system, dCas9-VPR, to induce expression of these pro-neuronal transcription factors to differentiate iPSC into neurons. We first transduced PBMC-iPSCs with lentiviruses carrying dCas9-VPR, followed by prolonged selection with puromycin to generate a stable iPSC line with dCas9-VPR (iPSC-VPR). Characterisation of iPSC-VPR using RT-PCR showed that the cells retain stable expression of the dCas9-VPR activator after prolonged culture (Figure 2I).

Using this iPSC-VPR system, we tested the feasibility of using specific sgRNA to transcriptionally activate *ASCL1, NEUROD1* and *NEUROG2*. We have previously tested sgRNAs for CRISPRa activation of *ASCL1* and *NEUROD1* ^*34*^, which target 181bp upstream and 158bp downstream of the transcriptional start site (TSS) respectively (Supplementary figure 1, Supplementary table 1). For *NEUROG2*, we designed a sgRNA targeting 135bp upstream of the TSS (Supplementary figure 1, Supplementary table 1). We introduced these sgRNAs into iPSC-VPR using lentiviruses and monitored the levels of gene activation using qPCR. Our results indicate that this CRISPRa approach can efficiently upregulate all three genes in iPSC, with high expression levels for *NEUROD1* and *NEUROG2* (∼1781 fold and ∼1832 fold increase respectively), followed by *ASCL1* (∼13 fold increase, Figure 2J). These results show that our CRISPRa iPSC system allows efficient activation of master transcription factors for neuronal specification.

Next, we tested the potential of using CRISPRa to direct iPSC differentiation into neurons, termed induced neurons (iN). To study how pro-neuronal transcription factors (*ASCL1, NEUROD1 or NEUROG2)* disrupt pluripotency and induce neuronal differentiation, we introduced the targeting sgRNA into iPSC-VPR while keeping the cells in stem cell medium. Our results showed that the sole activation of *ASCL1, NEUROD1* or *NEUROG2* are able to generate iN within 7 days (Figure 3). Notably, we observed the presence of cells with neuronal morphology as early as 4 days (Figure 3A-C). After 7 days, immunocytochemistry analysis showed that the iN expressed neuronal markers βIII-Tubulin (Figure 3D-G) and MAP2 (Supplementary Figure 2). In our testing, *NEUROD1* is the most efficient among the three transcription factors to generate iN characterised by long axons, therefore all subsequent iN generation experiments utilised *NEUROD1* activation. In summary, we have developed a CRISPRa iPSC platform that allows rapid neuronal differentiation to generate iN in 7 days.

**Figure 3.**
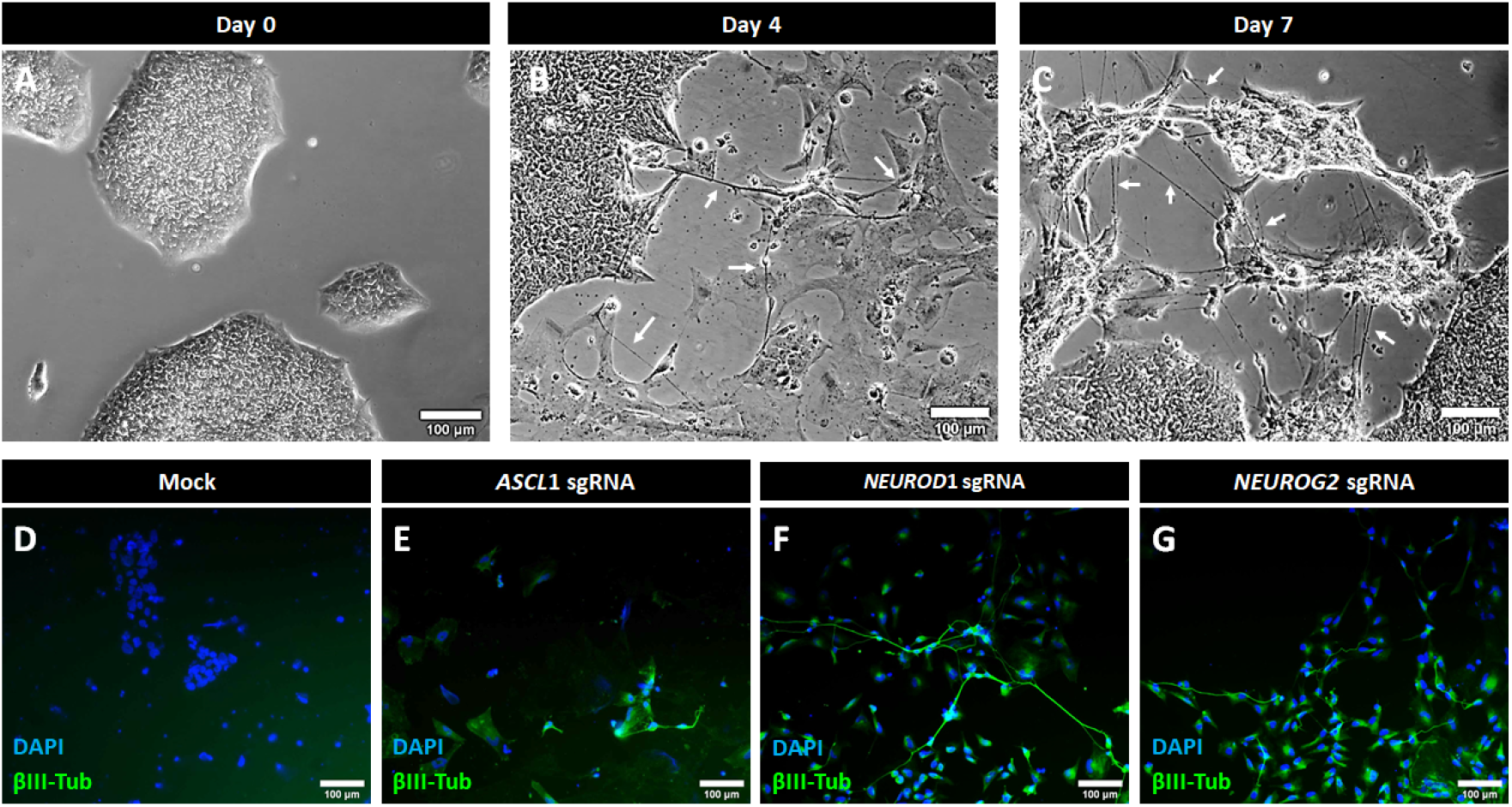
Using CRISPRa for transcription factor-directed neuronal differentiation. Representative morphology of *NEUROD1-*mediated differentiation to generate iN at A) day 0, B) day 4 and C) day 7. White arrows mark the derived iN with axons. Scale bar = 100 μm. D-G) Immunocytochemistry analysis of βIII-tubulin expression (green) and nuclei counterstain (blue) in iN generated by CRISPR activation of E) *ASCL1*, F) *NEUROD1*, G) *NEUROG2* and D) the relevant mock control. Scale bar = 100 μm.

### BCCs influence branching and neurite outgrowth

Surface topography can influence cellular processes such as morphology, proliferation and differentiation. To further improve our CRISPRa neuronal differentiation system, we investigated if BCCs with different topographies could improve the generation of iN using our CRISPRa iPSC platform. First, we assessed whether the BCCs could support stem cell growth in a feeder-free culture. iPSC-VPR were seeded on the three BCCs surfaces (BCC9, BCC13 and BCC16) coated with or without vitronectin, and cell attachment was assessed by DAPI staining. Our results showed that BCC with vitronectin coating provide better support for iPSC growth and a similar effect is observed in all three BCCs (Supplementary figure 3). On the other hand, iPSC growth is severely impacted on BCC surfaces without vitronectin coating. Thus, we proceeded to coat the BCCs with vitronectin for our neuronal differentiation protocol.

To evaluate the effect of different BCCs on neuronal fate induction, we seeded iPSC-VPR cells on BCCs and introduced a sgRNA to activate *NEUROD1* expression to generate iNs. Our results showed that all three BCCs surfaces are able to support iN generation to varying degrees (Figure 4A-D). Notably, our quantification results showed that all three BCC surfaces generate a significantly higher proportion of iN compared to flat glass control (11.11 ± 1.91%), with the highest iN proportion in BCC16 (33.56 ± 6.54%), while BCC9 and BCC13 generated a similar iN proportion (20.28% ± 3.91 and 20.68% ± 6.28 respectively, Figure 4E). These results suggest that utilisation of BCCs promote enrichment of neuronal culture, with the most prominent effect observed in the most hydrophilic and roughest surface (BCC16).

**Figure 4.**
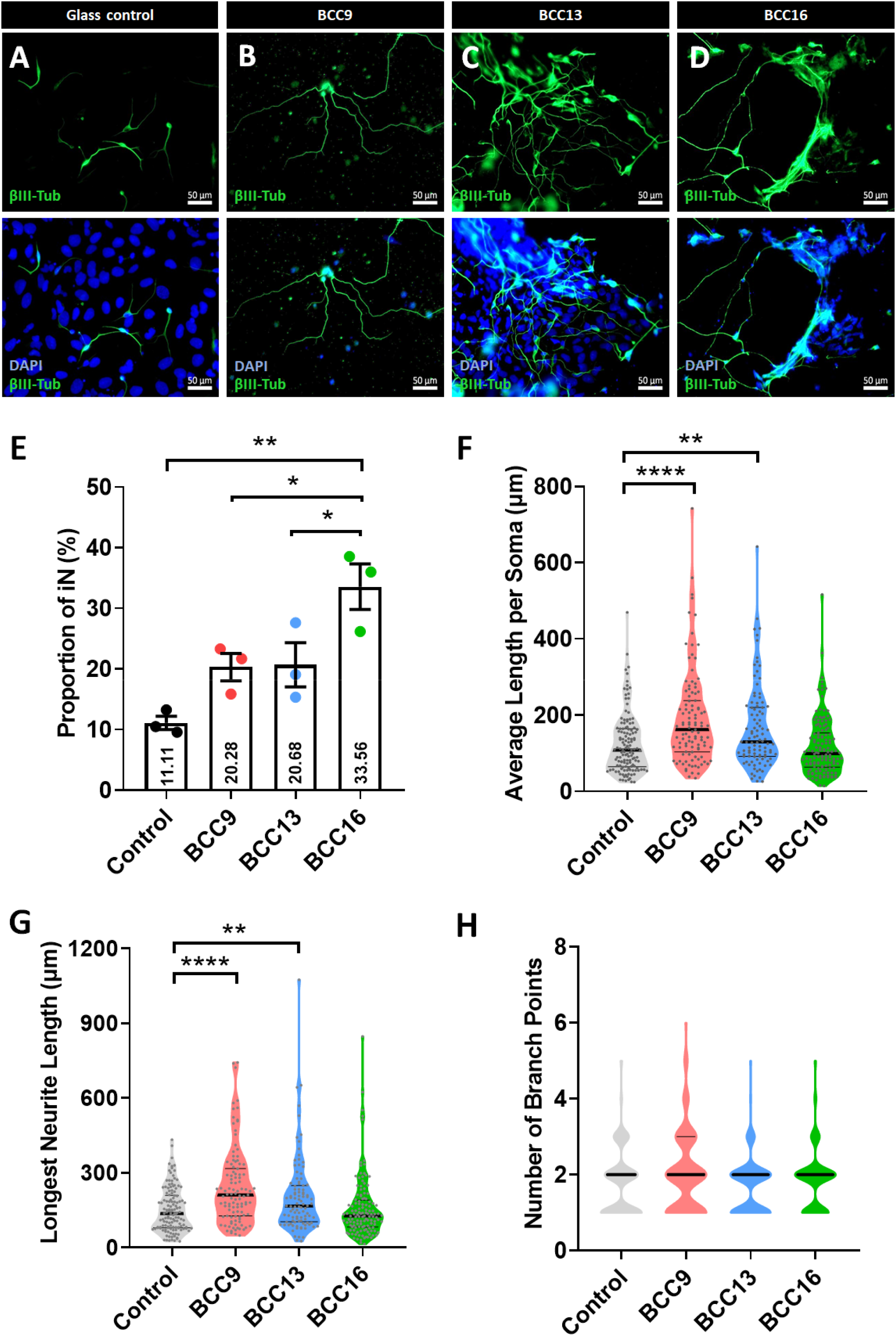
BCC surfaces improve iN generation, neurite outgrowth and ramification. Immunocytochemical analysis of iN derived on A) flat glass control, B) BCC9, C) BCC13 and D) BCC16, showing expression of βIII-Tubulin (top panel) and merged images with DAPI (bottom panel). E) Quantification of neuronal proportion in iN culture grown on flat glass control, BCC9, BCC13 and BCC16. n=3 biological repeats, error bars indicate SEM. *=p<0.05, ** = p<0.01. Violin plots showing quantification of the F) average neurite length, G) primary neurite length and H) neuronal branching in iN cultured on BCC and flat glass control. n= 3 biological repeats with a total of >100 neurons per condition.

Next, we investigated if the biophysical properties of BCCs could have an impact in neurite outgrowth and complexity, which are characteristics of functional mature neurons. Interestingly, our analysis of >100 iN cells showed that the different BCC topographies exert significant effects to several morphological features of the derived iN, including neurite length, outgrowth orientation and bendings on the neurites. We performed quantification analysis of the neurite outgrowth by comparing the average length of all the neurites per soma (Figure 4F), as well as the length of the longest primary neurite of the iNs (Figure 4G). Our results showed that the iNs produced on a flat glass control developed short neurites (averaged 124.4 μm per soma and 150.7 μm for primary neurite). In comparison, BCC9 and BCC13 promoted the development of longer neurites in iN, both in terms of the mean neurite length per soma (180.9 μm and 164.4 μm respectively, Figure 4F) and the mean primary neurite length (244.6 μm and 204.4 μm respectively Figure 4G). The longest primary neurite of iN was observed on BCC13 at 1074.83 μm, while BCC9 supported primary neurite growth as long as 742.8 μm (Figure 4G). In contrast, BCC16 only exerted a small effect on neurite outgrowth compared to flat glass control, with a mild increase to the longest neurite length but no significant changes to the overall average neurite length (Figure 4F-G).

In regard to neuronal ramification, our results showed that all BCCs support development of iNs with two neurites on average (Figure 4H). Interestingly, iNs on the BCC9 surface contained a bigger population with three or more neurites compared to the rest of the surfaces tested, suggesting that BCC9 can further improve on the neuronal complexity of iN. Collectively, these results showed that BCC9, that smoothest and most hydrophobic BCC tested, has the most prominent effect in promoting longer neurites and more complex branching. Our results provide supporting evidence that BCCs can be used together with our CRISPRa iPSC platform to generate iN with improved neurite outgrowth and ramification with more branching.

## Discussion

Pluripotent stem cells, such as iPSCs, have the exciting potential as an unlimited source of functional neurons. However, many neuronal differentiation protocols require months of differentiation and the addition of complex cocktails of growth factors and small molecules in multiple stages ^6,8,9^. In this study, we developed a simplified method for neuronal differentiation, taking advantage of a combinatorial use of CRISPRa to activate master transcription factor for neuronal specification and defined topographical control using BCCs.

The recent development of CRISPR technology has generated important tools that allow for endogenous gene regulation and epigenetic editing. Several CRISPRa systems have been developed using a nuclease deficient Cas9 (dCas9) coupled with transcriptional activators to induce potent gene expression, including the VPR ^25^, Suntag ^35^ and SAM systems ^36^. We have previously reported on the use of sgRNA expression cassettes for the dCas9-VPR system for multiplex gene activation ^34^. Building on this work, here we derived a new iPSC line using PBMC from a healthy donor and generated a stable line with constitutive expression of dCas9-VPR. Our CRISPRa iPSC platform provides an easy-to-use, efficient system to induce gene activation. Although CRISPRa has been utilised in neural progenitors to direct neuronal differentiation in a previous study ^37^, to our knowledge this study is the first to generate a CRISPRa iPSC platform with the dCas9-VPR system. Using this system, we showed that activation of key neural transcription factors, *ASCL1, NEUROD1* or *NEUROG2*, can drive iPSC to rapidly differentiate into neurons within 7 days. Notably, this neuronal differentiation process does not require addition of other proneural growth factors or small molecules, suggesting that the neural transcriptional networks activated by *ASCL1, NEUROD1* or *NEUROG2* are able to override the endogenous pluripotent transcriptional network and convert cell fate in iPSC. Our study created a CRISPRa iPSC platform that allows robust gene activation and expanded the previous findings that overexpression of *ASCL1, NEUROD1* or *NEUROG2* transgenes allow rapid neuronal differentiation ^7,10,11^. Notably, it is possible to use our CRISPRa iPSC platform for multiplex gene activation. Future studies that introduce additional transcription factors would allow us to explore the generation of specific neuronal subtypes, such as glutamatergic, GABAergic, dopaminergic, sensory and retinal neurons ^38^. Also, the implications of our CRISPRa iPSC system is not limited to differentiation into neurons; other transcription factors can be activated to direct differentiation into different lineages, such as hepatocytes ^39^, skeletal muscles ^40,41^ and pancreatic beta cells ^42^. These characteristics make our CRISPRa iPSC platform an attractive tool to generate *in vitro* models for multiple cell types.

To further improve on our CRISPRa neuronal differentiation process, we explored the effect of topography using various BCC surfaces. In recent years, the use of defined surface topography is emerging as a key strategy to control cell differentiation. In the present study, we showed that BCC surfaces with both 2 μm or 5 μm silica can promote enriched neuronal cultures. Our results expand the findings of previous studies that have used surface topography such as grantings, pillars and nanofiber scaffolds to support the generation of stem cell-derived neurons ^13,17–19^. In particular, some of the BCC surfaces are able to improve the generation of iNs with longer neurites and more complex branching, with the most prominent effect observed in BCC9 consisting of 5 μm silica and 100 nm carboxylated polystyrene, the smoothest and most hydrophobic surface among tested in this study. In contrast, the roughest and most hydrophilic BCC tested, BCC16 with 2 μm silica and 65 nm carboxylated polystyrene, has a strong effect on neuronal enrichment but no positive effect on neurite branching. This suggests that the fine tuning of silica and polystyrene size is critical to generate a specific topography that can improve neuronal differentiation and maturation. The length of the neurites from stem cell-derived neurons increases upon maturation, therefore neurite outgrowth and branching are indicative of maturation and functionality ^12,21^. In particular, BCCs with specific topographies could also be harnessed at specific stages of differentiation to fine-tune existing neuron differentiation protocols. For instance, Tan *et al*. found that grantings can aid early stages of neural differentiation, as they promote commitment into neural progenitors, whereas pillars increase branching and neuronal complexity that are more suitable for maturation ^19^. Given that some of the BCCs (BCC9 and BCC13) promoted the generation of longer neurites and/or more complex neurite branching, these surfaces could be adapted to the late stages of existing neuronal differentiation protocols to promote neuronal maturation.

Previous studies have demonstrated that cell-surface interactions can improve the kinetics of neuronal differentiation in pluripotent stem cells ^18^, as well as functionality of the derived neurons ^19,20^. Although the biochemical and genetic signals driving neuronal differentiation during development have been extensively studied, the precise mechanism of how biophysical cues affect neuronal differentiation remained largely unclear. For instance, topographies such as grantings can promote the upregulation of neuronal markers without the need of additional biochemical signals ^13,17^. One postulation is that the morphological changes induced by grantings could be, at least partially, responsible for favouring neuronal fate. Bridges and grooves promote culture alignment as well as elongation of cellular bodies to acquire a polarized morphology. This cytoplasm elongation is achieved by focal adhesion signals and cytoskeleton reorganization, which could in turn modify gene expression profiles in the nucleus ^13,43,44^. We speculate that the effect of BCCs on neuronal differentiation is exerted through a similar mechanism, where defined topographies caused morphological changes and cytoskeletal reorganisation in neurons, leading to downstream signaling that promote neurite elongation and branching. Future research that investigates the transcriptomic changes in iN cultured on BCCs would be interesting in addressing the downstream genetic signals modulated by topographical cues.

There are limitations to this study of the combinatorial use of BCCs and CRISPRa for neuronal differentiation. Although the current study determined that BCCs promote maturation through the formation of longer and more complex neurites, future study is needed to assess neuronal functionality, including action potential firing, synapse formation and neurotransmitter release which are beyond the scope of this study. Also, a limitation of the BCCs utilised in the present study was the reduced attachment rate compared to glass and polystyrene tissue culture plates. However, this can be resolved by plating more cells on BCCs to compensate for the reduced attachment and can be easily upscaled for neuronal differentiation. Alternatively, we could explore additional BCCs topographies that allow higher attachment ratios, while retaining the positive effects on neural differentiation shown by this study. Nevertheless, the highly ordered surface topographies of BCCs can be easily fabricated and upscaled to produce surfaces with different sizes and topographies. Thus, our simplistic approach for iPSC neuronal differentiation using biophysical cues and transcriptional activation can be easily upscaled, providing an exciting alternative to generate functional neurons *in vitro*.

In summary, this study presents a CRISPRa iPSC platform that can rapidly generate enriched neuronal culture without the need of complex biochemical signals and multiple differentiation stages. The fast acquisition of neuronal fate can be achieved by activation of *NEUROD1* and incorporation of BCC with defined topographies further improve neurite outgrowth and ramification. To our knowledge, this is the first report of a stable iPSC line with constitutive expression of the CRISPRa system, which represents a useful tool to activate transcription factors to drive *in vitro* differentiation into multiple cell types. Collectively, our results showed that the combinatorial approach of transcription factor-mediated differentiation with the defined surface of BCCs represents a promising strategy to generate enriched neuronal cultures with mature characteristics.

## Methods

### Cell culture

PBMC-iPSCs were cultured in feeder-free conditions in TeSR-E8 (Stem Cell Technologies) and maintained at 37 °C and 5% CO2. The culture plates were pre-coated with vitronectin (VTN-N; Thermo Fisher Scientific) for at least 1 h at room temperature, following the manufacturer’s instructions. For passaging, the cells were detached with ReLeSR (Stem Cell Technologies) following manufacturer’s instructions and transferred to a vitronectin-coated plate containing fresh medium with 10 µM of ROCK inhibitor Y227632 (Jomar Life Research).

### BCC fabrication and characterization

Monolayer BCCs were fabricated according to our previous method ^23^. In brief, three BCCs, BCC9 (5 μm Si particles and 100 nm carboxylated polystyrene), BCC13 (2 μm Si particles and 65 nm polystyrene), and BCC16 (2 μm Si particles and 65 nm carboxylated polystyrene) were selected for iPSC culture and differentiation. Glass was used as the control. The top-view structure of the BCCs was captured using field emission scanning electron microscopy (FE-SEM; ZEISS SUPRA 40 VP, Carl Zeiss, Germany) at 20 keV. The structure of BCCs as well as the surface property in terms of roughness and wettability were measured as we previously described ^22^.

### Donor blood collection and iPSC generation

Donor blood was collected from a 65 years old healthy male with informed consent ^26^, as approved by the Human Research Ethics Committees of the Royal Victorian Eye and Ear Hospital (11/1031H), in accordance with the guidelines from the National Health and Medical Research Council of Australia and in-line with the Declarations of Helsinki.

PBMC was isolated using CPT tubes following manufacturer’s instructions (BD Bioscience) and the erythroid progenitor cell was expanded using StemSpan media with Erythroid Expansion supplement (Stem Cell Technologies). Reprogramming of erythroid progenitor cells was performed using the Erythroid Progenitor Reprogramming Kit following manufacturer’s instructions (Stem Cell Technologies). Briefly, 10^6^ cells were nucleofected with the Epi5 reprogramming vectors, p53 and EBNA vectors, using the Human CD34+ cell nucleofector kit (Lonza) with program U-008. Following nucleofection, 3.3 x 10^5^ cells were plated down into one well of a 6-well plate pre-coated with matrigel in StemSpan media with Erythorid Expansion supplement. The media is switched to ReproTeSR medium 3 days later, with daily medium change for ∼21 days. Subsequently, iPSC colonies with morphology similar to human embryonic stem cells were manually picked and characterised as described previously ^27^.

### Lentivirus generation

One day prior transfection, 7×10^6^ HEK293FT cells were plated on a 10 cm dish and cultured with lentivirus packaging medium consisting of Opti-MEM supplemented with 5% FBS and 200 µM Sodium pyruvate (all from Thermo Fisher). The lentiviruses were generated by co-transfecting the HEK293FT cells using Lipofectamine 3000 (Thermo Fisher) with the following plasmids: lenti-EF1a-dCas9-VPR-Puro plasmid (gift from Kristen Brennand; Addgene, #99373) or the sgRNA expression cassette lentiGuide-Puro (gift from Feng Zhang; Addgene, #52963), with the three packaging vectors pMDLg/pRRE (Addgene, #12251), pRSV-Rev (Addgene, #12253) pMD2.G (Addgene, #12259). The medium was replaced with fresh medium six hours post-transfection, and the viruses were collected at 48 and 72 hours post-transfection. The collected viruses were filtered (0.45 µm filter, Sartorius) and concentrated using PEG-it overnight at 4°C (SBI Integrated Sciences). The virus titre was calculated using the ELISA-based Lenti-X™ p24 Rapid Titer Kit (Takara Bio) following the manufacturer’s instructions.

### Generation of iPSC-VPR cell line

PBMC-iPSCs were transduced overnight with lentiviruses that co-express dCas9-VPR and the puromycin resistant gene (EF1a-dCas9-VPR-P2A-Puro) in the presence of 8 µg/mL of polybrene (Sigma-Aldrich). Two days post-transduction, the cells were selected with 500 ng/mL of puromycin (Thermo Fisher Scientific) to establish a stable iPSC-VPR cell line.

### Embryoid body formation assay

*In vitro* differentiation of iPSC was performed using embryoid bodies (EB) formation as previously described ^28,29^. Briefly, the EBs were formed by seeding the iPSCs into a low attachment plate and maintained in suspension culture for 11 days. Subsequently, the EBs were plated on a gelatin coated plate and further differentiated for 18 days. The samples were processed for immunocytochemistry to assess expression of the three germ layer markers smooth muscle actin (SMA), alpha-fetoprotein (AFP) and βIII-Tubulin.

### Karyotype analysis

Genomic DNA of PBMC-iPSC was extracted using the Wizard Genomic DNA Purification kit (Promega) following manufacturer’s instructions. Karyotype analysis was performed using the single nucleotide polymorphism assay iCS-digital PSC test (Stem Genomics).

### Transcription factor-mediated neuronal differentiation using CRISPRa

iPSC-VPR cells were seeded on coverslips with binary colloidal crystals (BCCs) pre-coated with vitronectin in a 24 well plate, and allowed to grow for 2 days. On day 0, the cells were transduced with lentiviruses with sgRNA for activating *NEUROD1, ASCL1* or *NEUROG2* (MOI = 5). Lentiviral transduction was performed overnight in TeRS-E8 medium with 8 µg/mL of polybrene (Sigma-Aldrich). On day 1, the virus was removed and replaced with fresh TeRS-E8. From day 3 to day 7, the cells were cultured with Neurobasal medium supplemented with B27 (Thermo Fisher). On day 7 the cells were fixed and immunostained for neural markers.

### Immunocytochemistry

Standard immunocytochemistry procedures were carried out as previously described ^30^. Briefly, samples were fixed in 4% paraformaldehyde, followed by permeabilization in 0.1% Triton X-100. The samples were blocked with 10% Goat Serum (Sigma-Aldrich) and incubated at 4 °C overnight with antibodies against βIII-Tubulin (Millipore, MAB1637), MAP2 (Thermo Fisher, MA512826), SMA (R&D systems, MAB1420) or AFP (Millipore, ST1673). Subsequently, the samples were stained with Alexa Fluor 488 secondary antibodies (Millipore) for 1 hour at room temperature, followed by nuclear counterstain with DAPI (Sigma-Aldrich). Images were taken using an Axio Imager.M2 microscope (Zeiss) and analysed with the ZEN 3.2 Blue edition software.

### Analysis of neuronal induction

To quantify the neurons, images were taken using the tiles tool of ZEN Blue edition (ZEISS), each image consisted of 16 individual picture frames (∼ 1.67 x 1.25 mm). For each surface, a total of four images from random regions were taken in three independent differentiation replicates. To quantify the proportion of iN, βIII-Tubulin positive neurons were counted using the Cell Counter plugin of ImageJ and DAPI staining was used to quantify total cell number.

For neurite analysis, the neurite outgrowth of the iN was marked by βIII-Tubulin staining and quantified using the ImageJ plugin NeuronJ ^31^. For each sample, individual neurons from five random fields were analysed by tracing and measuring their neurites. Three independent neuronal induction replicates were carried out and a total of at least 100 neurons were measured for each surface condition.

### qPCR analysis

Total RNA samples were extracted and processed for DNaseI treatment using the Illustra RNAspin kit (GE Healthcare). cDNA synthesis was performed using the High-Capacity cDNA Reverse Transcription Kit with RNase Inhibitor (Thermo Fisher), following the manufacturer’s instruction. qPCR was performed using the TaqMan gene expression assay (Thermo Fisher Scientific) with the following probes: NEUROD1 (Hs00159598_m1), ASCL1 (Hs00269932_m1), NEUROG2 (Hs00702774_s1) and the housekeeping control ACTB (Hs99999903_m1). The TaqMan assay was processed using the StepOne plus (Thermo Fisher).

### Statistical test

Unpaired two-tailed Student’s t-test was performed for neuron quantification and one-way ANOVA test was performed for CRISPR activation using Graphpad Prism. p<0.05 was used to assess statistical significance.

## Additional Information

The authors declare no conflict of interest.

## Data Availability

The authors declare that data supporting the findings of this study are available within the paper and its supplementary information files.

## Acknowledgement

We would like to sincerely thank Aaron Liu for technical support for this study. This research was supported by the University of Melbourne (RCBW) and the Centre for Eye Research Australia (RCBW). DU and RL are supported by the Melbourne Research Scholarship from the University of Melbourne. The Centre for Eye Research Australia receives operational infrastructure support from the Victorian Government. PYW is supported by the Ministry of Science and Technology of China (2019YFE0113000); the National Natural and Science Foundation of China (31870988); the Science, Technology, and Innovation Commission of Shenzhen Municipality (20180921173048123; ZDSYS20190902093409851).

## Author contribution

RCBW and PYW designed the experiments; DUC, RHCL, JL, KL and SSCH conducted the experiments; DUC, RHCL, JL, KL, SSCH, AWH, PYW and RCBW analysed the data; AH, PYW, RCBW provided funding to this work; DUC, RHCL, PYW and RCBW wrote the manuscript. All authors approved the manuscript.

## Supplementary figures

**Supplementary Figure 1.**
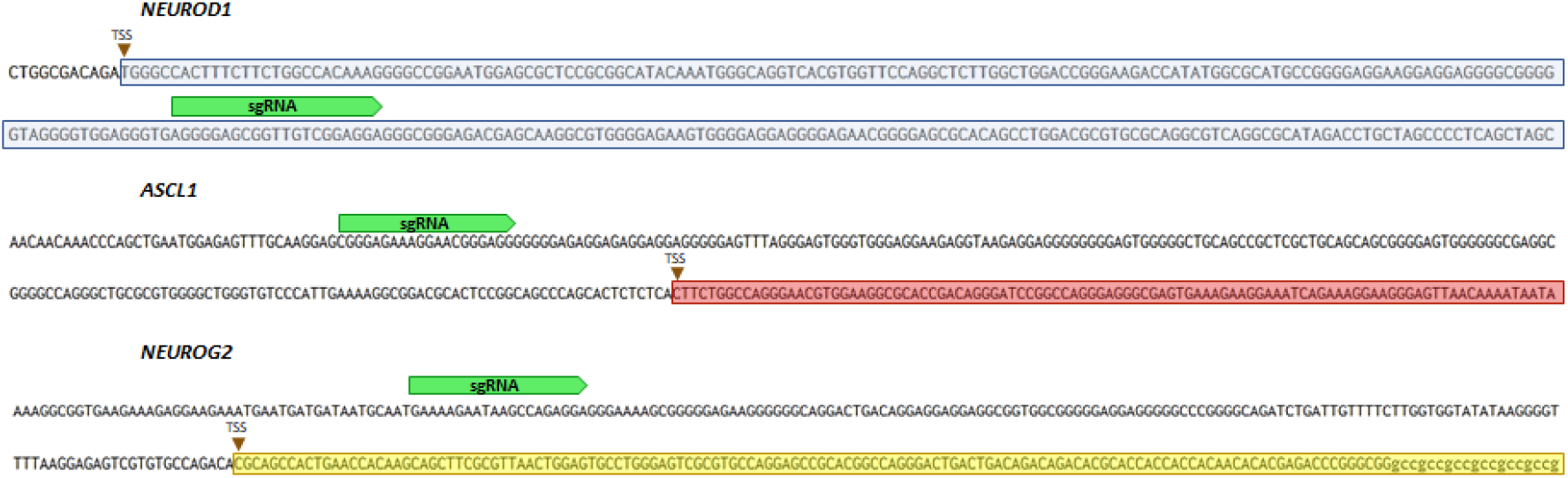
Diagram of sgRNA target areas (green) near the transcription start site (TSS) of the human genes *NEUROD1* (blue), *ASCL1* (red) and *NEUROG2* (yellow).

**Supplementary Figure 2.**
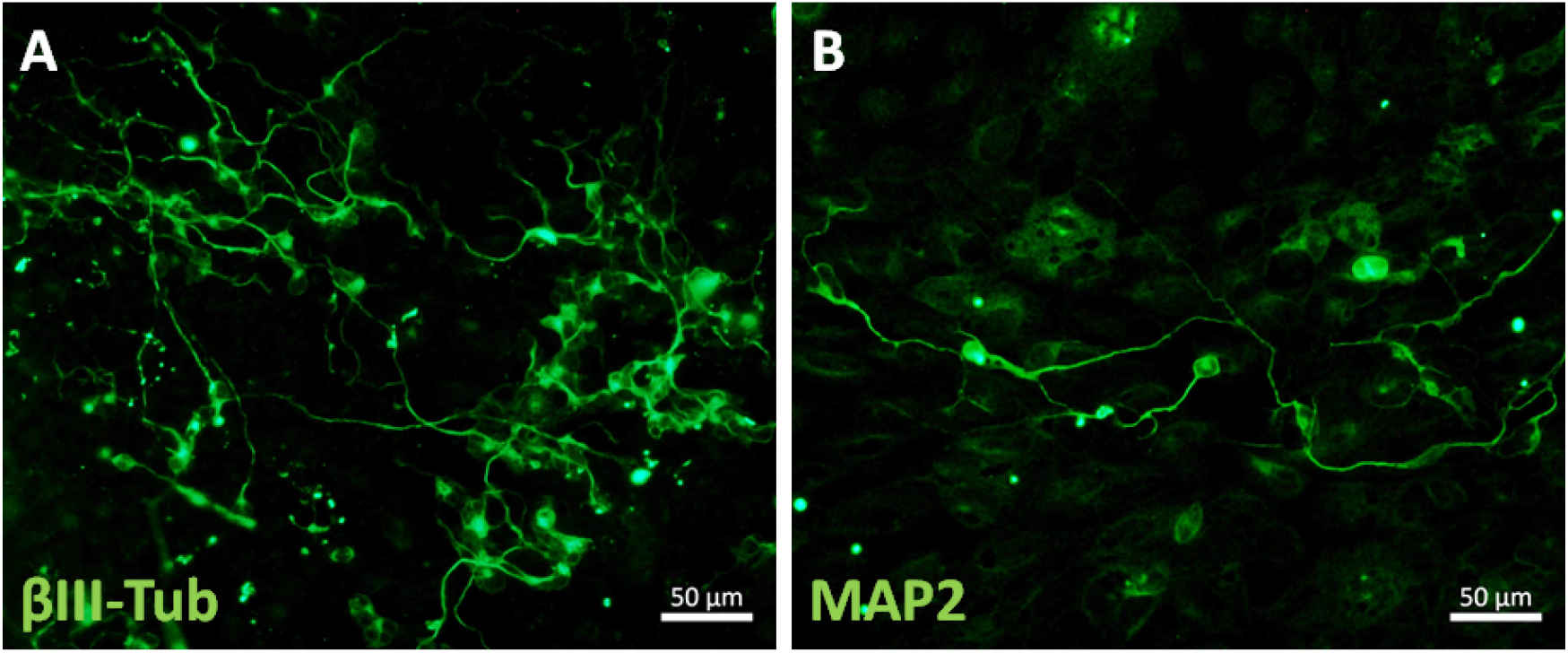
*NEUROD1-*mediated differentiation to generate iN with positive expression of neuronal markers A) βIII-tubulin and B) MAP2. Scale bar = 50 μm.

**Supplementary Figure 3.**
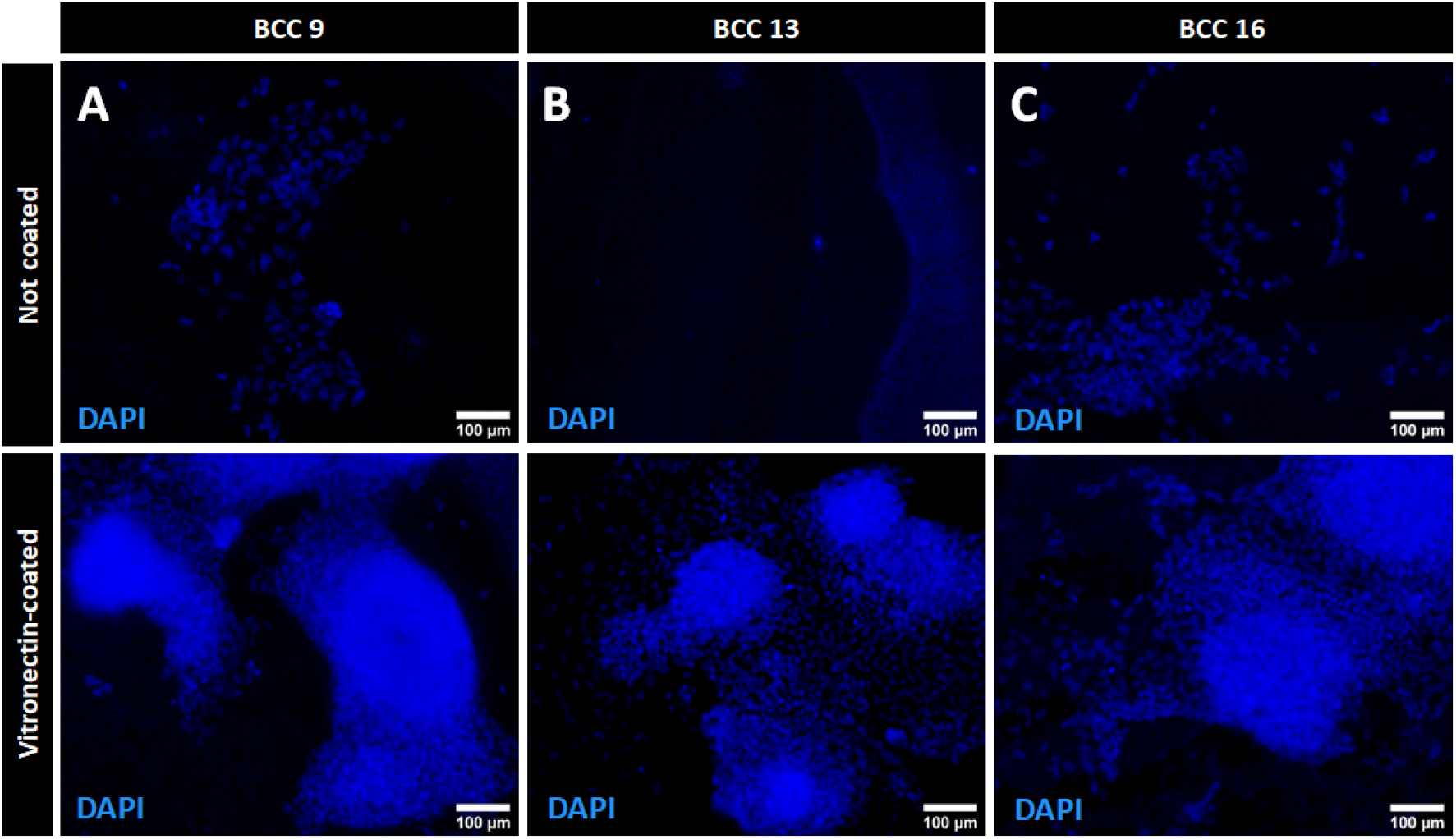
Assessment of iPSC attachment on A) BCC9, B) BCC13 and C) BCC16. Representative picture of DAPI counterstain of iPSC cultured on A-C) non-coated BCCs or A’-C’) BCCs coated with vitronectin. Scale bar = 100 μm.

**Supplementary table 1:**
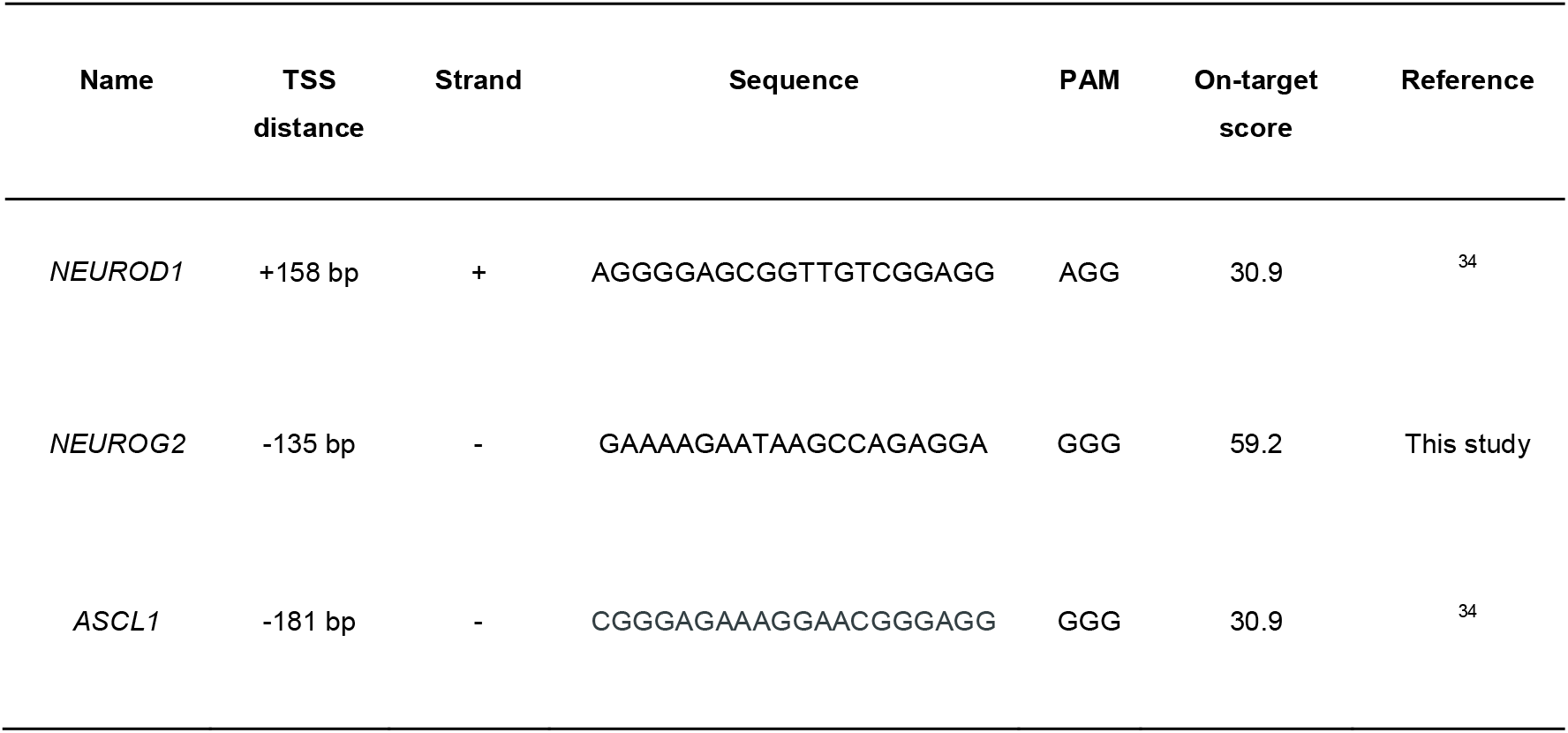
Information of sgRNAs used in this study. TSS distance is based on the transcription start site defined by Ensembl. On-target score is based on ^45^.

## Notes

### Competing Interest Statement

The authors have declared no competing interest.

